# Packing-Driven Mechanotransduction: local crowding overrides adhesion and stiffness cues for YAP Activation in Cellular Collectives

**DOI:** 10.1101/2025.10.03.680214

**Authors:** Valeriia Grudtsyna, Vinay Swaminathan, Amin Doostmohammadi

## Abstract

The regulation of mechanotransduction is crucial for various cellular processes, including stem cell differentiation, wound healing, and cancer progression. While the activation of mechanotransduction has been extensively studied in single cells, it remains unclear whether similar mechanisms extend to mechanotransduction in multicellular collectives. Here, by focusing on Yes-associated protein (YAP), known as the master regulator of mechanotransduction, we reveal that the local packing fraction of cells acts as the primary determinant of YAP activation in cell collectives. We further show that local packing fraction modulates the isotropic stress landscape, with sparse regions experiencing large stress fluctuations and dense regions displaying stress equilibration. Remarkably, this packing fraction-dependent regulation persists even under conditions of disrupted force transmission through cell-cell and cell-substrate adhesion, suggesting a robust and conserved relation between YAP activation and local packing fraction in cell collectives. In particular, we show that local packing fraction-dependent activation of YAP in cell collectives is independent of substrate stiffness, E-cadherin expression, and myosin contractility, in stark contrast to YAP activation in single cells. Our results thus offer a new perspective on mechanotransduction, highlighting the critical role of local packing fraction of cells in dictating YAP dynamics within multicellular contexts. These insights have significant implications for tissue engineering and understanding tumour microenvironments, where cellular heterogeneity often drives functional outcomes.

## Introduction

Cells in tissues constantly integrate biochemical and mechanical cues from their microenvironment, with mechanical cues being highly localized due to processes such as division, extrusion, and intercellular interactions, including pushing and pulling [1]. Mechanotransduction enables cells to interpret these inputs, influencing their behaviour and maintaining tissue homeostasis through feedback loops [2, 3]. One of the key pathways involved in mechanotransduction is the Hippo signalling pathway, a conserved regulatory cascade influenced by environmental cues. This pathway plays a central role in controlling cell proliferation, differentiation, and survival by modulating gene expression [4]. YAP, a key effector of this pathway, shuttles between the cytoplasm and nucleus depending on its phosphorylation state, with mechanosensitive importins mediating its nuclear translocation via nuclear pore complexes [5]. Other upstream regulators of the Hippo signalling pathway include G-protein coupled receptors (GPCRs) [6], metabolic signals [7], and cellular energy status [8]. These regulators integrate various extracellular and intracellular signals to modulate YAP activity and ensure appropriate cellular responses.

What makes YAP particularly fascinating is its regulation by both biochemical and mechanical pathways. Since the discovery of the mechanical route [9], it has become evident that forces, force transmission, and mechanical cues can be crucial and even override biochemical pathways in regulating YAP activity. This dual regulation underscores the importance of YAP in various cellular processes and its potential as a therapeutic target.

Among the most important mechanical pathways influencing YAP are actomyosin contractility, substrate stiffness, cell-cell adhesion, and nuclear mechanics. Actomyosin contractility, which drives mechanoreciprocal force generation [10, 11], is a key player in mechanotransduction, with evidence showing that YAP promotes actin polymerization to regulate cell rigidity and, in turn, relies on actin integrity for nuclear localization [9, 12, 13]. This highlights a feedback loop between YAP activity and cytoskeletal dynamics. Similarly, studies suggest that YAP tends to shuttle into the nucleus on stiff 2D substrates [9, 12, 14, 15], although conflicting reports indicate no consistent correlation between substrate stiffness and the nuclear-to-cytoplasmic YAP ratio, whether in single cells [16], 2D monolayers [17], or 3D acini [18]. This suggests that force transmission mechanisms beyond substrate stiffness alone may be at play.

In addition to the cell-substrate coupling, cell-cell adhesion also plays a central role in mechanical signalling and YAP regulation. Intercellular force transmission and contact inhibition of proliferation are highly dependent on the E-cadherin/catenin complex [19, 20]. This complex is frequently discussed in the context of YAP activation, defined as the YAP nuclear-to-cytoplasmic (n/c) ratio [21, 22, 23, 24] and nuclear to cytoplasmic shuttling of YAP has been observed in E-cadherin–negative parental breast cancer cell line upon insertion of E-cadherin [25].

The nucleus itself represents another important regulatory hub in mechanotransduction. Nuclear compression, often mediated by forces transmitted through the actin cytoskeleton and cell-substrate adhesions, flattens the nucleus and reduces mechanical restrictions at nuclear pores, enabling YAP translocation into the nucleus [14]. The interplay between cell and nuclear volume, as well as molecular crowding, can further impact YAP dynamics [26, 27]. However, how nuclear shape and mechanics influence YAP localization in multicellular systems remains underexplored.

While most of our understanding of YAP regulation comes from studies on single cells [9, 16], important biological processes and diseases are often multicellular effects. Epithelial cells function collectively to maintain the integrity of the tissue layer, making isolated cell behaviour less relevant. Furthermore, monolayers are highly heterogeneous in terms of other properties, such as cell shape [28], cell stiffness [29], and metabolism [30], meaning that global averaging of cellular properties can obscure critical local variations. Understanding how upstream regulators of the Hippo pathway, such as substrate stiffness, E-cadherin, and actomyosin contractility, impact YAP activation in the multicellular context requires consideration of these local heterogeneities.

In this study, we demonstrate that within cellular collectives, the local packing fraction of cells is the primary determinant of YAP activation. Remarkably, we show that this regulation persists even when force transmission through cell-cell and cell-substrate adhesion is disrupted. Our findings demonstrate that packing-driven YAP activation in cell collectives is independent of substrate stiffness, E-cadherin expression, and myosin contractility, which contrasts with the mechanisms observed in single cells. These results highlight the critical role of local packing fraction of cells in regulating YAP activation, independent of force transmission mediated by cell-cell and cell-substrate adhesion, or nuclear morphology. We propose that local packing fraction of cells is a relevant and descriptive metric in multicellular systems, significantly influencing cellular processes, morphology, and expression, which are often obscured when globally averaged.

## Results

### Characterization of MDCK monolayer: relation between YAP activation and local packing fraction

MDCK cells were cultured at varying seeding densities on fibronectin-coated glass-bottom dishes and allowed to proliferate overnight. Following incubation, cells were immunostained using an anti-YAP antibody to assess YAP localization. Notably, YAP localization exhibited considerable heterogeneity across the MDCK monolayer (Figure 1a). At low global cell densities (Figure 1a, left), substantial variability in YAP localization was observed, with distinct clusters of cells displaying elevated nuclear YAP, while others exhibited predominantly cytoplasmic YAP. Cells with similar YAP localization were frequently found in proximity to one another; however, occasionally, cells deviated from their immediate neighbours.

**Fig. 1.**
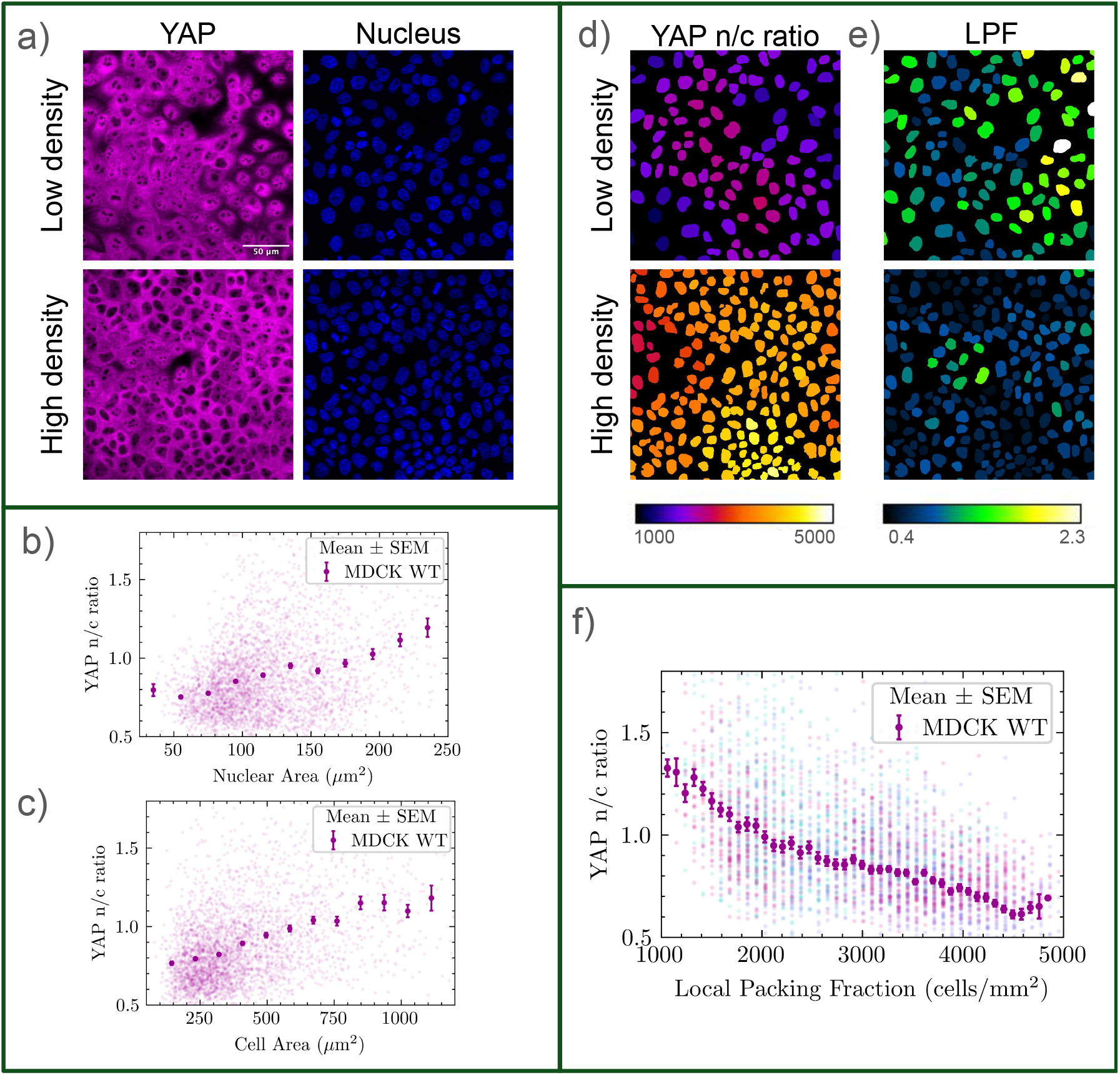
Characterization of MDCK monolayer: relation between YAP activation and local packing fraction. a) MDCK monolayer at low and high global density, immunostained with YAP and dyed with Hoechst. These representative images illustrate the spatial heterogeneity of YAP n/c localization at various global densities. b) YAP nuclear/cytoplasmic ratio (n/c) as a function of nuclear area (3380 cells combined from 3 experiments). Binned data is presented as mean and standard error of the mean. c) YAP nuclear/cytoplasmic ratio as a function of estimated cell area (3380 cells combined from 3 experiments). Binned data is presented as mean and standard error of the mean. d) Cell nuclei segmentation colour-coded as the YAP n/c ratio for the same cell patch as in a). e) Cell nuclei segmentation colour-coded as the local packing fraction for the same cell patch as in a). f) YAP n/c ratio versus local packing fraction (3324 cells combined from 3 experiments). The radius of the local packing fraction window is 60 *µ*m. Binned data is presented as mean and standard error of the mean.

As global cell density increased (Figure 1a, right) and the average cell area decreased, YAP localization predominantly shifted to the cytoplasm, resulting in a reduction of spatial variation in YAP distribution across the monolayer. This is in agreement with previous single-cell studies, which have consistently reported a correlation between the basal cell area and the YAP nuclear-to-cytoplasmic (n/c) ratio [9, 31, 32, 33]. To further examine this relationship, we quantified YAP n/c ratios in relation to the nuclear area (Figure 1b,c). Consistent with prior observations, a general trend emerged wherein more extensively spread cells, characterized by larger projected nuclear areas, exhibited increased YAP activity. However, substantial variance in YAP n/c ratios was evident, indicating heterogeneity in cellular responses.

Cells with similar YAP n/c ratios tended to cluster within the monolayer (Figure 1d). A potential factor influencing this spatial organization is the local packing fraction, quantified as the number of cells within a predefined radius (60 *µ*m) (Figure 1e). Analysis of the relationship between YAP n/c ratio and the local packing fraction revealed a consistent trend (Figure 1f), demonstrating higher variance in YAP activation at low packing fractions. This was further underscored by standard linear regression and the Spearman correlation analysis (Supporting Figure 6a). Additionally, PCA analysis was performed to highlight the YAP - local packing fraction correlation (Supporting Figure 6b). This observation parallels the heterogeneity observed at low global densities, suggesting that sparse conditions may contribute to greater variability in YAP-dependent cellular responses.

These results suggest that local cell density plays a role in regulating YAP activity within MDCK monolayers. However, YAP is also known to be responsive to mechanical cues, including force transmission through cell-cell junctions and intracellular cytoskeletal tension. Actin cytoskeletons of the neighbouring epithelial cells are connected to each other via adherens junctions, which enable the transmission of forces between cells. At the same time, actin is connected to the ECM via focal adhesions, enabling transmission of traction forces to the underlying substrate as well as sensing of the external forces and substrate stiffness. We thus asked how the observed relation between YAP activation and local packing fraction is affected by force transmission mediated by cell-cell and cell-substrate interactions.

### Role of cell-substrate and cell-cell force transmission in YAP activation is overridden by local packing fraction of cells

We began by altering the force transmission between the cells and the ECM by changing the substrate stiffness [34]. Substrate stiffness-dependent YAP activation has been one of the cornerstone findings in the study of mechanotransduction. Numerous studies have demonstrated that YAP becomes more nuclear on substrates of increasing stiffness, highlighting the critical role of mechanical cues from the extracellular matrix in regulating cellular behaviour [9, 35, 14, 15].

Cells were cultured on commercially available elastic substrates with stiffnesses of 1.5 and 15 kPa, precoated with fibronectin, and compared to a standard glass control. These stiffness values were selected to cover an order of magnitude in stiffness and span a range over which YAP is known to transition toward cytoplasmic localization as stiffness decreases [36]. The level of actin stressfibers is low on 15 kPa and is further decreased on 1.5 kPa (Figure 2a). Representative images of YAP immunofluorescence at low and high global densities (Figure 2b) revealed no obvious differences in YAP localization between the glass control and the softer substrates. Strikingly, the relation of measured YAP n/c ratio versus local packing fraction (Figure 2c) showed overlapping trends across more than order of magnitude variation in substrate stiffness, suggesting a conserved relation between local packing fraction and mechanotransduction activation. Having observed no effect from varying substrate stiffness, we then turned into examining the impact of altering force transmission through cell-cell adheison.

**Fig. 2.**
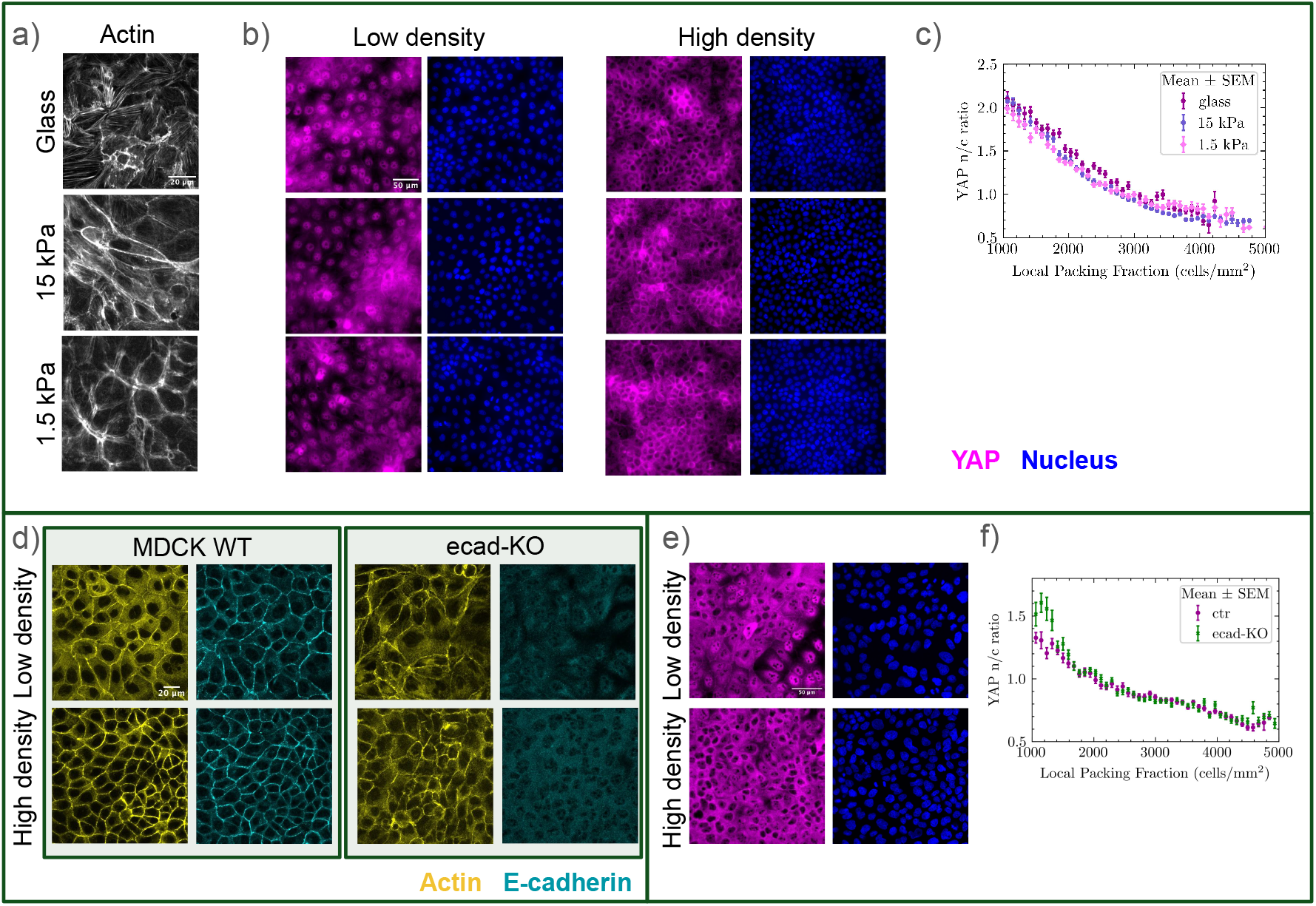
Role of cell-substrate and cell-cell force transmission in YAP activation is overridden by local packing fraction of cells. a) Representative images of cortical actin stress fibers on glass, 15 kPa and 1.5 kPa. F-actin is stained with Phalloidin. There is a decrease in stress fibers and altered architecture as the substrate stiffness drops. b) Representative images showing YAP staining and the nuclear channel of WT MDCK monolayers at high and low global density, plated on glass, 15 kPa and 1.5 kPa. The images are taken using wide-field epifluorescence. c) YAP n/c ratio versus local packing fraction for the three substrate stiffnesses (6322, 6099, 5855 cells on glass, 15 kPa, 1.5 kPa. Combined data from 3 experiments). d) Representative images of WT and E-cadherin KO MDCK cells at high and low global density, confirming that E-cadherin is excluded from cell junctions, but F-actin remains intact at both junctional and basal levels. e) Representative images showing E-cadherin KO MDCK cells high and low global density immunostained with YAP and dyed with Hoechst. f) YAP n/c ratio versus local packing fraction (3324 WT and 3688 E-cadherin KO cells from 3 experiments). Binned data is presented as mean and standard error of the mean.

Mechanical force transmission is crucial in mechanotransduction within cell collectives as it enables cells to sense and respond to their physical environment [37]. E-cadherin plays a key role in this process by mediating cell-cell adhesion, which is essential for coordinating cellular responses and maintaining tissue integrity [38, 39]. Previous studies have highlighted the significance of E-cadherin expression in YAP activation [25, 21, 23, 24].

To investigate the role of E-cadherin, we utilized an MDCK E-cadherin knockout (KO) cell line, in which E-cadherin is excluded from cell junctions (Figure 2d). Notably, F-actin remains intact at both the junctional and basal levels. Similar to wild-type MDCK cells, E-cadherin KO cells exhibit patches of varying local packing fractions at both seeding densities, and YAP expression patterns are comparable to those in wild-type cells (Figure 2e).

Based on previous reports on the defining impact of E-cadherin on YAP activation [25, 21, 23, 24], we expected that in the absence of E-cadherin, the relation between YAP n/c ratio and local packing fraction would change substantially. Unexpectedly, however, we found the relationship between the YAP n/c ratio and local packing fraction nearly indistinguishable between the two cell types (Figure 2f). As with variations in substrate stiffness, control and E-cadherin KO monolayers showed substantial overlap in YAP localization across the full range of local packing fractions. Only a small subset of highly nuclear YAP E-cadherin KO cells appeared at very low packing fractions, where the local environment effectively corresponds to the individual cell limit. This indicates that E-cadherin is not required for the packing fraction-dependent regulation of YAP in the multicellular context.

The results presented so far challenge the previously understood role of substrate stiffness and E-cadherin in YAP activation and underscore the overriding influence of local packing fraction. These results emphasize the importance of studying YAP activation in the context of multicellular collectives rather than single cells, suggesting the existence of a robust and conserved relation between YAP activation and local packing fraction. While single-cell studies have provided valuable insights into mechanotransduction, the collective behaviour of cells within tissues introduces additional layers of complexity and regulation.

### Actin integrity, but not myosin contractility, is essential for maintaining the robust relationship between YAP and local packing fraction

Given that force transmission did not affect the YAP - local packing fraction relationship, we next investigated the role of force generation in YAP activation. The actin cytoskeleton, composed of relatively stiff fibers, interacts with myosin to generate and resist forces [40]. The influential role of actin on YAP activation has been earlier established in single cell systems or as an averaged response at sparse/dense conditions [9, 12]).

To elucidate the role of actomyosin contractility in the interplay between YAP and local packing fraction, we first inhibited nonmuscle myosin II with Blebbistatin (50 *µ*M for 3 hours). Blebbistatin stops myosin II from generating force by docking between the ATP-binding site and actin-binding interface, thus locking it in a non-force-generating state and preventing the power stroke [41]. The treatment resulted in a relaxed cortical actin network with less defined stress fibers (Figure 3b). Interestingly, however, and in congruence with the observed behaviour for force transmission through cell-cell and cell-substrate adhesions, the packing-driven YAP activation remained robust against Blebbistatin treatment (Figure 3b). The relationship between YAP activation and local packing fraction was comparable to the control case for packing fractions >2000 and showed lower activations for packing fractions <2000(Figure 3c). This suggests that pure actomyosin contractility can enhance YAP activation in low density conditions, where the local environment is effectively corresponding to the individual cell limit. Importantly, however, as with the effect of force transmission, the relationship between YAP activation and local packing fraction remains independent of myosin-inhibition in dense monolayers.

**Fig. 3.**
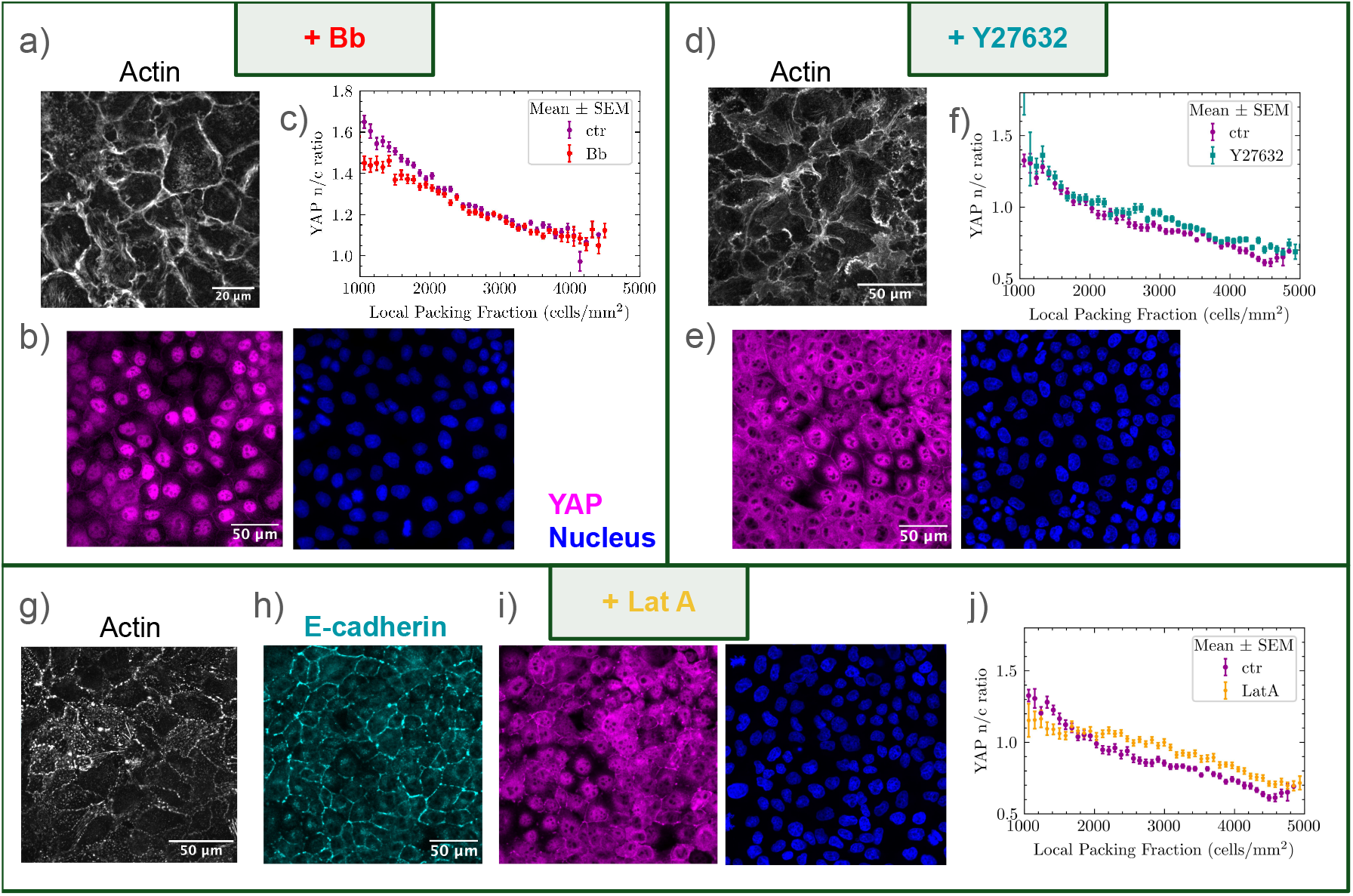
Actin integrity, but not myosin contractility, is essential for maintaining the robust relationship between YAP and local packing fraction. a) Representative images of cortical F-actin stressfibers in Blebbistatin treated cells, showing a relaxed cortical actin network with less defined stress fibers. b) Representative images of Blebbistatin treated cells immunostained for YAP and dyed with Hoechst. c) YAP n/c ratio versus local packing fraction (6858 control and 6128 Blebbistatin treated cells from 3 experiments). d) Representative images of basal F-actin stressfibers in Y-27632 treated cells, showing a less visible stress fibers and disrupted cortical actin patterns. e) Representative images of Y-27632 treated cells immunostained for YAP and dyed with Hoechst. f) YAP n/c ratio versus local packing fraction (3324 control and 3653 Y-27632 treated cells from 3 experiments). g) Representative images of basal F-actin stressfibers in cells treated with Latrunculin A. LatA caused disassembly of actin stressfibers and aggregates of cortical actin. h) Representative images of E-cadherin immunostaining in cells treated with Latrunculin A. i) Representative images of YAP immunostaining and the nuclear channel. in cells treated with Latrunculin A. j) YAP n/c ratio versus local packing fraction (3324 control and 3404 LatA treated cells from 3 experiments). Binned data is presented as mean and standard error of the mean.

We then asked whether interfering with actin filament organization could affect the conserved YAP-local packing relation. To this end, we first used Y-27632 (25 *µ*M for 4 hours), a compound that inhibits Rho-associated kinases 1 and 2 (ROCKs). These kinases regulate various cellular processes related to the actin cytoskeleton, such as controlling actin filament organization and cellular contractility by phosphorylating several downstream targets, including myosin light chain phosphatase [42]. This treatment caused less visible stress fibers and disrupted cortical actin patterns (Figure 3d). Despite these visible changes in actin architecture, the pattern of YAP activation remained similar to that of untreated cells (Figure 3e). Here, the relationship between YAP activation and local packing fraction instead showed that cells treated with Y27632 have elevated nuclear YAP in cells at packing fractions >2000 (Figure 3f). This suggests the suppressive role of ROCKs in YAP activation in dense, multicellular systems.

Building on these observations, we then explored whether more extreme perturbations—such as dismantling the actin cytoskeleton, a central structural and signalling component—might break this conserved relationship. To test this, we disrupted actin integrity using Latrunculin A (LatA), which prevents actin polymerization and promotes depolymerization [43, 44]. Cells treated with LatA (0.5 µM for 30 minutes) showed rapid disassembly of actin filaments, formation of actin aggregates at cell-cell junctions and less consistent E-cadherin expression along the junctions (Figure 3g,h). LatA treatment significantly altered the visual appearance of YAP as well as the YAP - packing fraction relationship (Figure 3i,j). At low cell packing fractions (*<*2000 cells/mm^2^), YAP n/c ratio was markedly reduced compared to controls. Conversely, at higher packing fractions (*>*2000 cells/mm^2^), the n/c ratio was elevated relative to untreated cells. Overall, the response curve became flattened, indicating that dismantling the actin cytoskeleton disrupts the packing fraction-dependent modulation of YAP localization. Nevertheless, YAP n/c ratios remained partially responsive to packing fraction, suggesting that other mechanisms also contribute to this regulation.

These results reveal that the integrity of the actin cytoskeleton is essential for maintaining the robust relationship between YAP localization and local packing fraction. While other perturbations, ranging from changes in force transmission to myosin contractility, failed to break this relationship across the whole range, dismantling the actin cytoskeleton represents a critical threshold where this robustness is lost. This finding highlights the central role of actin in integrating mechanical and biochemical signals that govern YAP activity, offering new insights into how cellular architecture shapes mechanotransduction.

### Nuclear shape properties are governed by local packing fraction, but influence YAP activation only at low local packing fractions

The cell nucleus has been widely implicated in mechanotransduction, with properties such as the nuclear volume [16] and flattening [14] proposed as potential regulators of YAP localization. Previous studies have explored various nuclear shape properties and their correlation to YAP activation at the single-cell level. For example, nuclear flattening induced by force enhances nuclear YAP import in mouse embryonic fibroblasts [14] and increased nuclear volume is associated with reduced YAP nuclear localization in NIH-3T3 cells [16]. A recent study [45] further correlates higher YAP activation with increased nuclear solidity index, defined as the ratio of the nucleus’s area to the area of its convex hull.

We investigated whether nuclear shape contributes to the robust relationship between YAP localization and local packing fraction in our system. As we found no clear relation between the local packing fraction and the two-dimensional nuclear aspect ratio (Supporting Figure 7a), we turned towards the three-dimensional segmentation. Reconstructed confocal stacks of cell nuclei were segmented using Cellpose (Figure 4a) and analysed for nuclear volume *V*, height *h*, (Figure 4b,c), flatness (defined as nuclear length *L* divided by height *h*) (Figure 4d,e), shape factor (defined as *S/V* ^2*/*3^) (Supporting Figure 7b) and surface area *S* (Supporting Figure 7c), across MDCK monolayers under four conditions: wild type (WT), E-cadherin knockout (KO), ROCK inhibition (Y27632), and Latrunculin A (LatA) treatment.

**Fig. 4.**
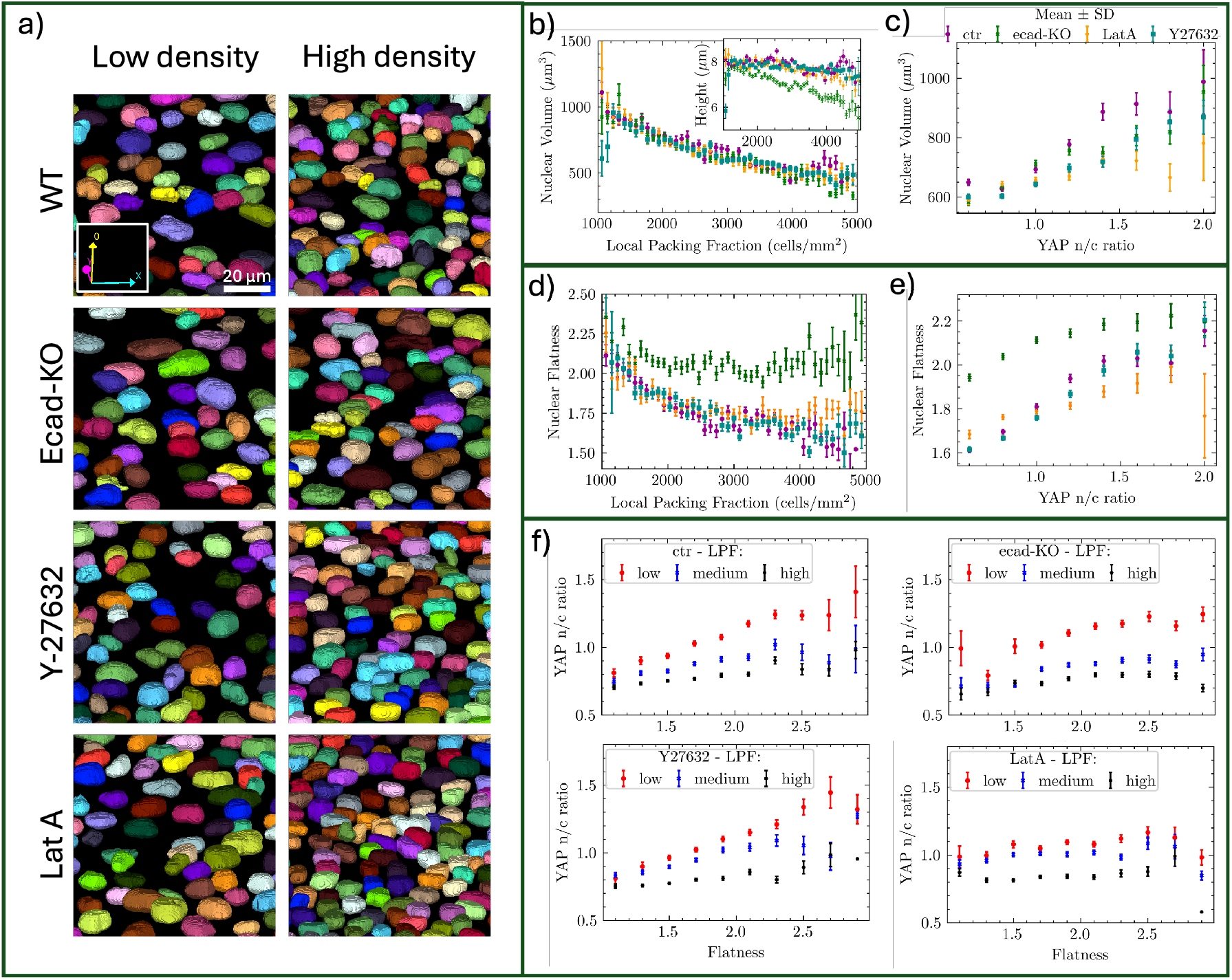
Nuclear shape properties are governed by local packing fraction and possibly influence YAP activation at low local packing fractions. a) Nuclear segmentation of WT, E-cadherin KO cells and of WT cells treated with Y-27632 and LatA at high and low density. The scale bar is 20 *µ*m. b-e) Plots of conditions shown in a). 3324, 3688, 3653 and 3404 cells of WT, E-cadherin KO, Y-27632 and LatA treated WT cells. b) Nuclear volume versus local packing fraction. The inset plot is showing nuclear volume versus local packing fraction. c) Nuclear volume versus YAP n/c ratio. d) Nuclear flatness (defined as nuclear length/height) versus local packing fraction. e) Nuclear flatness versus YAP n/c ratio. f) Same data as in b-e), binned by local packing fraction into low (lower than 2300 cells/mm^2^, medium (2300 - 3270 cells/mm^2^) and high (above 3270 cells/mm^2^). YAP n/c ratio is plotted versus the nuclear flatness for WT, E-cadherin KO, Y-27632 and LatA treated WT cells. Binned data is presented as mean and standard error of the mean.

While nuclear height showed a small decline at high packing fractions (Figure 4b), nuclear volume decreased systematically with increasing cell packing fraction across all conditions, approximately halving as packing fraction increased from 1000 to 5000 cells/mm^2^. YAP n/c ratio and nuclear volume showed a clear correlation, with higher YAP activation in cells with larger nuclei (Figure 4c). At high local packing fractions, nuclei became less flat (Figure 4d), possibly because cells were more confined and nuclei approached a more spherical shape. Flatter nuclei correlated positively with YAP n/c ratio at all conditions (Figure 4e).

Despite these pronounced changes in nuclear shape properties across conditions, YAP localization remained robustly regulated by local packing fraction. For example, while LatA treatment disrupted the conserved YAP-packing fraction relationship, nuclear volume, surface area, and shape factor remained packing fraction-dependent and unchanged compared to other conditions. Similarly, E-cadherin KO cells displayed significantly altered nuclear shape, yet their YAP localization trends with packing fraction were indistinguishable from WT cells. These findings strongly suggest that nuclear shape properties do not drive the robust relationship between YAP localization and packing fraction.

To test whether the nuclear shape has different impact at different packing fractions, we binned the data into four groups by packing fraction. We first tested nuclear flatness and volume. When plotting nuclear flatness against the YAP nuclear-to-cytoplasmic (n/c) ratio (Figure 4f), we found that nuclear flatness had the greatest influence on the YAP n/c ratio in sparsely packed cells, with its impact diminishing as cell density increased. However, under LatA treatment, nuclear flatness did not significantly affect YAP regulation, as indicated by the flat curves. Similarly, nuclear volume, when plotted against the YAP n/c ratio (Supporting Figure 7d), showed a weaker correlation, suggesting that these nuclear shape factors were not the primary determinants of YAP activation.

Given these results, we turned our attention to DNA density, as it can influence nuclear mechanics and chromatin organization, which are key factors in gene expression regulation. To investigate whether DNA density could be a potential regulator of YAP activation, we measured the mean Hoechst intensity as a proxy for DNA density [14] and plotted it similarly for binned packing fractions (Supporting Figure 7e). Our analysis revealed that in control MDCK and E-cadherin knockout (KO) cells, there was a clear dependence of YAP localization on DNA density, particularly at low packing fractions. This indicates that DNA density significantly impacts YAP activation in these conditions. However, this trend was less pronounced in cells treated with Y-27632 and LatA. The flatter curves in these treated cells suggest that the disruption of cytoskeletal integrity diminishes the influence of DNA density on YAP activation. These findings highlight the complex interplay between nuclear characteristics and YAP regulation, emphasizing the importance of considering multiple factors in understanding YAP dynamics.

Together, these results challenge the notion that nuclear mechanical factors govern YAP localization. Instead, they put forward a picture in which at low packing fractions, nuclear morphology could potentially have a role in regulating YAP, but that the importance of nuclear morphology diminishes as the cells are experiencing a more packed environment. Nuclear deformation might be a consequence of the local packing fraction, rather than a driver by itself. While untangling the complex interplay between nuclear characteristics and YAP regulation remains beyond the focus of the current work, our results suggest that nuclear morphology can serve as a secondary regulator of YAP signalling, following local packing fraction.

### Cells at lower local packing fractions experience tensile stresses and strong stress fluctuations

Cells exert and experience varying levels of mechanical stress depending on their local environment and density [46][47]. To link local packing fraction with the mechanical environment of cells, we quantified isotropic stress distributions using Bayesian Inversion Stress Microscopy (BISM) [46][48], which infers intercellular stress from the balance between cell–substrate traction and intercellular forces (Fig. 5a).

**Fig. 5.**
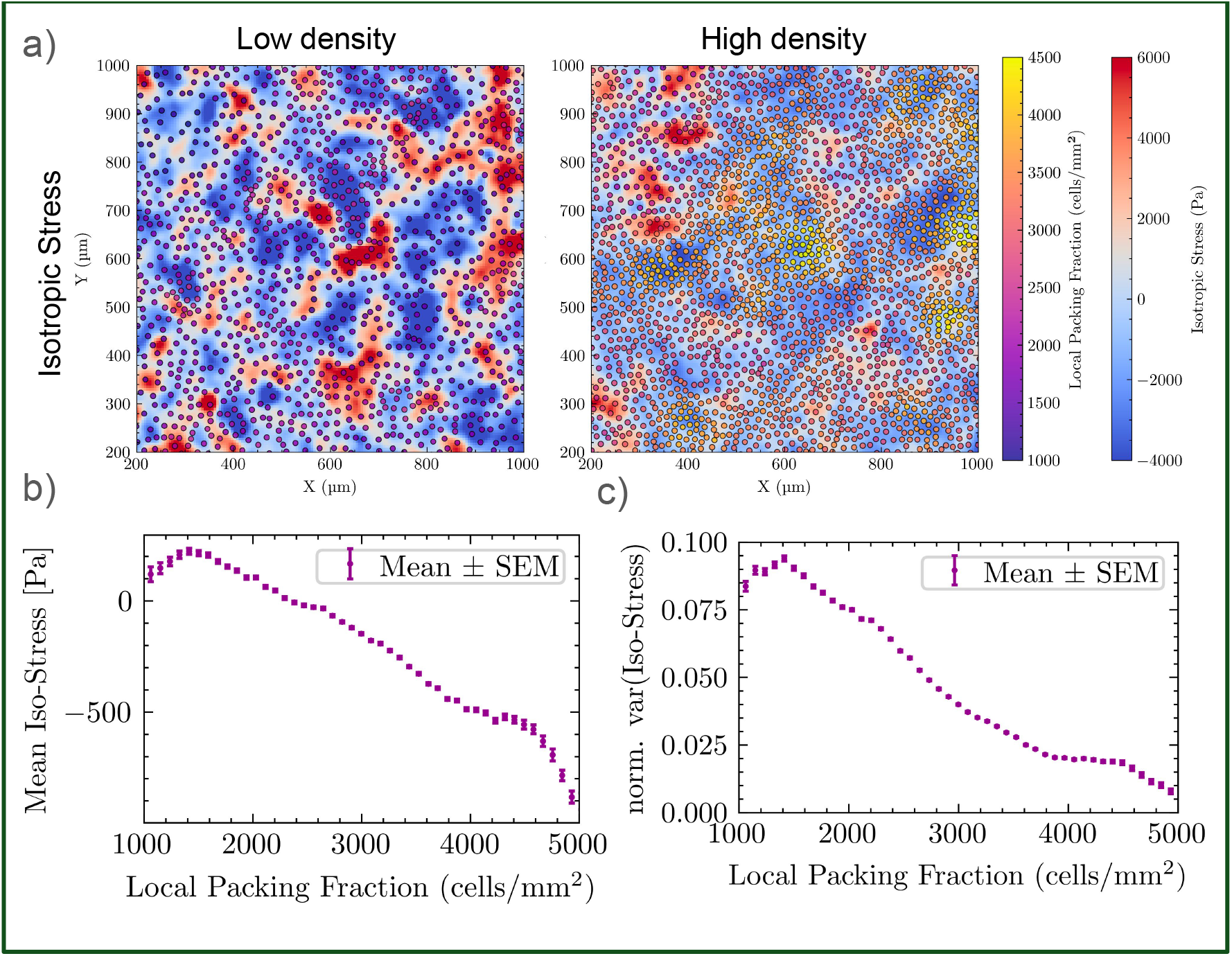
Isotropic stress. a) Isotropic stress field at low and high global cell density. Red regions are corresponding to tension and blue to compression. Cell centroids are scattered on top and colour-coded based on the local packing fraction. b) Mean isotropic stress as a function of local packing fraction. The mean isotropic stress is quantified for the same region as the local packing fraction (radius of the window is 60 *µ*m). 597077 cells are in total analysed. Binned data is presented as mean and standard error of the mean. c) Normalized variance of isotropic stress as a function of local packing fraction. The variance is quantified for the same region as the local packing fraction (radius of the window is 60 *µ*m). 597077 cells are in total analysed. Binned data is presented as mean and standard error of the mean.

**Fig. 6.**
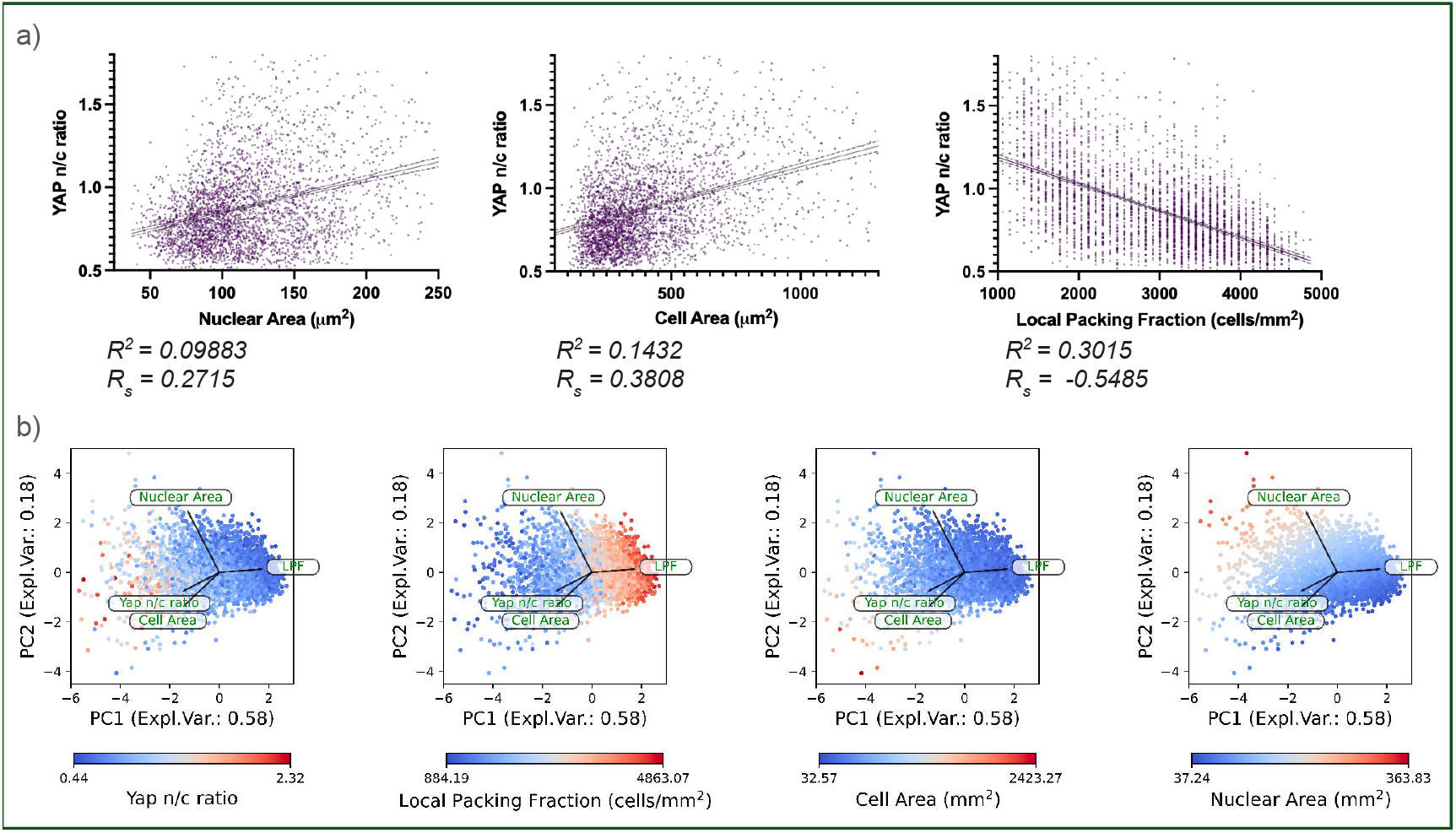
Linear regression analysis. a) YAP nuclear/cytoplasmic ratio (n/c) as a function of nuclear area, cell area and local packing fraction (3380 cells combined from 3 experiments). Simple linear regression with 95% confidence intervals. Underneath the plots are the corresponding *R*^2^ values together with respective *R*_*s*_ Spearman correlation. b) PCA analysis applied on YAP n/c ratio, cell area, nuclear are and local packing fraction. PC1 explains 58% of variance and PC2 explains 18% of variance. Each plot is colour-coded based on each of the four chosen properties.

**Fig. 7.**
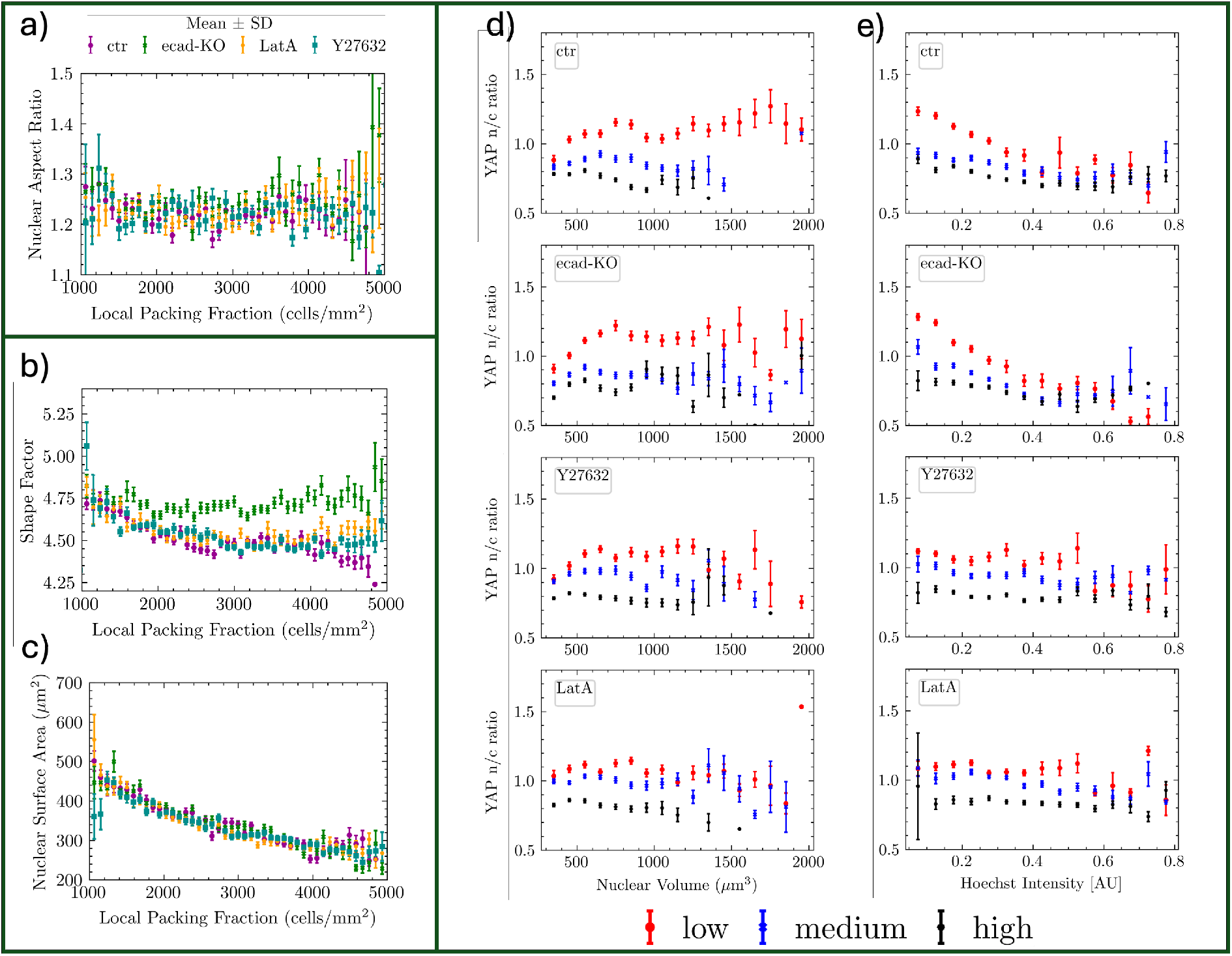
Other measured nuclear shape properties and Hoechst mean intensity (proxy for DNA density) plotted for following conditions: WT, E-cadherin KO, Y-27632 and LatA. a-d) 3324, 3688, 3653 and 3404 cells of WT, E-cadherin KO, Y-27632 and LatA treated WT cells. a) Nuclear aspect ratio (a 2D property) plotted versus local packing fraction. b) Nuclear shape factor as a function of local packing fraction. c) Nuclear surface area as a function of local packing fraction. d) Same data as in a-c), binned by local packing fraction into low (lower than 2300 cells/mm^2^, medium (2300 - 3270 cells/mm^2^) and high (above 3270 cells/mm^2^). YAP n/c ratio is plotted versus the volume flatness for WT, E-cadherin KO, Y-27632 and LatA treated WT cells. e) Same data as in a-c), binned by local packing fraction into low (lower than 2300 cells/mm^2^, medium (2300 - 3270 cells/mm^2^) and high (above 3270 cells/mm^2^). YAP n/c ratio is plotted versus Hoechst mean intensity for WT, E-cadherin KO, Y-27632 and LatA treated WT cells. Binned data is presented as mean and standard error of the mean.

At low local densities, cells on average experience tensile isotropic stress, whereas at higher densities the mean stress progressively shifts toward compressive values, consistent with crowding-induced confinement (Fig. 5b). In addition to this shift in the mean, the variance of isotropic stress shows a marked dependence on local packing fraction. Sparse regions display large stress fluctuations, reflecting high mechanical susceptibility, while in dense regions the fluctuations decrease nearly threefold and reach a low, saturating level (Fig. 5c).

These results suggest that local crowding transforms the stress landscape from one dominated by variable tensile and compressive loads to one characterized by mechanical equilibration. In sparse neighborhoods, poor coupling between cells leaves individual cells exposed to strong and fluctuating stresses, conditions that promote nuclear YAP/TAZ localization. In dense neighborhoods, force transmission and crowding buffer stress fluctuations, reducing susceptibility and favoring cytoplasmic YAP/TAZ. We therefore propose a Packing–Susceptibility Conjecture: local packing fraction governs YAP/TAZ dynamics by modulating a cell’s exposure to stress fluctuations, with nuclear YAP/TAZ favored under high susceptibility and cytoplasmic localization favored under buffered conditions. We suggest that examining how tissues regulate such susceptibility represents a promising avenue for future studies.

## Discussion

Yes-associated protein (YAP) is a pivotal regulator of mechanotransduction, influencing critical cellular processes such as stem cell differentiation, wound healing, and cancer progression. Understanding the regulatory mechanisms of YAP, particularly the interplay between local packing fraction of cells and mechanical cues, is essential for advancing tissue engineering and elucidating tumour microenvironments. Within epithelial monolayers, local crowding and mechanical forces create distinct “microenvironments” that dictate YAP activity, driving cellular behaviour heterogeneity at the tissue scale.

Here we elucidated the key regulators of YAP in an epithelial monolayer model, with a particular focus on determining whether possible robust determinants of YAP activation exist in the multicellular context. Surprisingly, by comparing substrates of significantly different stiffness (15 and 1.5 kPa), our results established that substrate stiffness does not significantly influence YAP localization within the tested range. This observation challenges the established notion that substrate stiffness is a determining factor for YAP activation and suggests that local packing fraction of cells can over-ride the effects of substrate stiffness. Future investigations could explore cellular responses to even softer substrates or substrates of different compositions to further elucidate this phenomenon.

The role of E-cadherin in packing fraction-dependent YAP regulation was also examined. The results indicate that E-cadherin is not required for packing fraction-dependent YAP regulation in this experimental setup. This finding is surprising given the well-documented importance of E-cadherin in YAP activation. The minimal role of E-cadherin in this context may be attributed to the presence of other mechanosensitive proteins, such as other cadherins, catenins, and integrins. In E-cadherin knockout (KO) cells, the nucleus is overall flatter, with a larger reduction in height over the packing fraction range. These cells maintain a constant shape factor, suggesting that their nuclear deformation compensates for the loss of junctional tension. Despite the altered nuclear shape, E-cadherin loss does not affect the packing fraction-dependent YAP regulation, underscoring the overriding influence of local packing fraction of cells.

Similarly, Blebbistatin treatment showed that nonmuscle myosin II-mediated contractility does not appear to be a dominant factor in regulating YAP at higher packing fractions, while it does affect YAP activation at lower packing fractions. In contrast, ROCK-mediated effects suppress YAP activation in MDCK cells at higher densities, as evidenced by an elevated response when the Y-27632 inhibitor is added. In this vein, more extreme perturbation of the integrity of actin skeleton through using Latrunculin A (LatA) treatment resulted in the flattening and attenuation of the packing fraction-dependent YAP response. This suggests that the integrity of actin filaments plays a critical role in regulating YAP shuttling across packing fraction ranges. At low packing fractions, disrupted actin integrity likely impairs force generation and mechanotransduction, resulting in reduced nuclear YAP localization. At higher packing fractions, LatA-induced cytoskeletal and junctional disruption may reduce cell-cell junction tension, softening the monolayer and enhancing nuclear YAP accumulation. The attenuated—but still detectable—sensitivity of YAP to packing fraction, even upon dismantling actin integrity, suggests that local packing fraction remains a key determinant. However, LatA likely interferes with other regulatory processes, such as cell junction integrity, which might indirectly affect YAP localization.

Nuclear volume, surface area, and 2D area are consistent across all conditions (control, E-cadherin KO, LatA, and Y-27632), suggesting that nuclear compaction is a conserved response to increasing local packing fraction of cells. Additionally, our results suggest that nuclear flatness, volume and DNA density are more influential at low packing fractions. This consistency further supports the notion that local packing fraction of cells is a fundamental regulator of YAP activation.

Together, our findings highlight the critical role of local packing fraction of cells as a primary regulator of YAP activation in epithelial monolayers, independent of substrate stiffness, E-cadherin expression, and myosin contractility. Despite variations in these mechanical and biochemical factors, local crowding remains a dominant factor driving YAP dynamics. Consistent with this, our stress analysis using Bayesian Inversion Stress Microscopy revealed that cells at low local packing fractions experience tensile stresses with strong fluctuations, while dense regions shift toward compressive stresses with reduced variability. This transition in the mechanical landscape provides a plausible mechanism for packing fraction dependent YAP regulation, where high stress fluctuations favor nuclear YAP/TAZ and buffered conditions promote cytoplasmic localization.

These results suggest that in any multicellular context, considering local packing fraction of cells is crucial, as it has a profound impact on cellular behaviour, morphology, gene expression, and mechanical state. This is especially important when studying tissues, where heterogeneity often governs cellular functions and responses. These insights have significant implications for tissue engineering and understanding the complex microenvironments of tumours, where cellular heterogeneity plays a pivotal role in disease progression and therapeutic resistance.

## Methods and Materials

### Cell lines

MDCK-II (ATCC CCL-34) and MDCK-II E-cadherin knockout (clone B6P6, as described in [48]) cell lines were generously provided by the Benoît Ladoux group. Cells were cultured in Dulbecco’s Modified Eagle Medium (DMEM, low glucose, GlutaMAX™Supplement, pyruvate) supplemented with 10% Fetal Bovine Serum (FBS; Gibco) and 1% Penicillin-Streptomycin (Gibco) at 37°C in a humidified atmosphere containing 5% CO_2_. All cell lines were routinely tested for mycoplasma contamination.

### Experimental procedure

Samples were prepared on 35 mm glass-bottom dishes No. 1.5 (Mattek) or equivalent glass-bottom dishes (FD35-100; FluoroDish). For substrate stiffness experiments 35 mm dish with 15 and 1.5 kPa Elastically Supported Surface (Ibidi) were used. Prior the cell seading, glass-bottom dishes were coated with 10 *µ*g/ml human fibronectin (F0895, Sigma-Aldrich) and elastic Ibidi substrates were functionalized with fibronectin according to the supporting manual.

Cells were ether seeded at two different packing fractions (1000 and 2000 cells/mm2) or spread unevenly over the substrate to create a wide range of local packing fractions and incubated over night. After ca 24 hrs, cells formed monolayers with distinct global densities and were fixed, stained with phalloidin and hoechst as well as immunostained for YAP, E-cadherin. A perturbation drug was added prior to fixation in stated concentration and for stated amount of time: Blebbistatin Racemic (203389; Sigma-Aldrich) 50 *µ*M for 3 hrs, Y-27632 (72304; StemCell) 25 *µ*M for 4 hrs, Latrunculin A (L5163; Sigma-Aldrich) 0.5 *µ*M for 30 min. Fixed and stained monolayers were mounted using non-solidifying vectashield, to avoid shrinkage in height.

### Immunostaining

Cells were fixed with 4% paraformaldehyde in PBS for 10 min at RT, permeabilized with 0.2% Triton TX-100 in PBS for 5 min, and blocked with 1% BSA in PBS for 45 min. Cells were immunostained with primary antibodies (mouse-anti YAP monoclonal IgG2a antibody (sc-101199; Santa Cruz biotechnology) (1:400) or the combination of rabbit-anti YAP1 polyclonal IgG antibody (13584-1-AP; ProteinTech) (1:250) and rat-anti E-cadherin Monoclonal IgG2a Antibody (ECCD-2) (131900 F; ThermoFisher Scientific) (1:250)) in 1% BSA in PBS and incubated at 4°C overnight or for 3 hours at RT (room temperature). Cells were washed three times with PBS and incubated for 2 h at RT in secondary antibodies (goat anti-mouse Alexa 568-conjugated IgG antibody (11004) or the combination of Goat anti-Rabbit 568 568-conjugated IgG antibody (A11011) and Goat anti-Rat 568 568-conjugated IgG antibody (A11006) (Life Technologies Corporation)), Alexa Fluor 488 Phalloidin (Invitrogen) (1:500) and nuclear dye Hoechst 350 (33342 Solution; Thermo Fisher Scientific) (1:4,000). Samples were washed and mounted in VECTASHIELD® PLUS ((H-1900), Vector Laboratories).

### Imaging

Samples were imaged on a spinning disk confocal microscope (Nikon) with a 40x water objective. Z-stacks with 1 *µ*m step-size were taken to capture the whole cell volume. Substrate stiffness samples and samples treated with Blebbistatin were imaged in wide-field mode using a Cell Observer Spinning Disk (Zeiss) with a 40x water objective. A single frame image was taken (focusing on the nuclear channel). Basal actin fibers were imaged using the spinning disk confocal.

### Image analysis

In-house Python pipelines were written to perform image analysis. Image processing package pyclesperanto was extensively used [49].

YAP n/c ratio: the central z-plane of the nuclear channel was segmented using Cellpose [50], to define the nuclear portion. These segments were dilated using the py-clesperanto package and a slightly dilated nuclear segmentation was subtracted, to define the cytoplasmic portion. This cytosolic portion is doughnut shaped around the nucleus and thus excludes the immediate transition area that connects the nucleus and the cytoplasmic as well as cell edges, where the cell is thinner. The YAP n/c ratio is calculated by dividing the mean intensity of YAP in the nuclear portion by the mean intensity of YAP in the cytoplasmic portion. The YAP intensities used in this calculation are from the central slice, mid plane of the cell. For the substrate stiffness experiment, a single plane wide-field image was taken and was used for the YAP n/c ratio analysis. This image was first background subtracted using top-hat.

For cell area estimation, the segmented nuclei were expanded by applying pyclesperantos voronoi labelling.

The local packing fraction of cells was calculated by applying py-clesperanto’s proximal neighbour count map onto the nuclear segmentation. The radius was set to 60 *µ*m (except 30 *µ*m for the window size comparison in the supplementary). The nuclear shape properties: the nuclear channel is segmented in 3D using Cellpose [51]. The shape properties are calculated using py-clesperanto’s statistics of labelled pixels.

### Statistical analysis

Statistical analyses were performed using GraphPad Prism version 10. Linear regression analysis was used to assess relationships between YAP n/c ratio and local packing fraction and results are reported as the coefficient of determination (*R*^2^). Spearman correlation analysis was performed for nonparametric associations, and results are reported as Spearman’s correlation coefficient (*R*_*s*_). PCA analysis was performed using Scikit-learn [52].

#### A. Bayesian Inversion Stress Microscopy

PDMS substrates were prepared by mixing CY 52-276 components at ratios of 6:5 (3 kPa) or 1:1 (15 kPa) (previously described in [53]), spreading the mixture in Petri dishes, and curing at 80 °C for 2 h. Gels were silanized with 10% (3-aminopropyl)triethoxysilane (APTES, Sigma-Aldrich) in ethanol for 10–15 min, rinsed, dried, and coated with carboxylate-modified fluorescent microspheres (1:500, ultra-sonicated, 5 min). After rinsing and drying, substrates were coated with 10 µg/ml human plasma fibronectin (in PBS) for 1–2 h at 37 °C.

Cells were seeded onto bead-coated PDMS substrates and cultured overnight to form nearly confluent monolayers. Live imaging was performed on a Nikon ECLIPSE Ti microscope equipped with an Okolab chamber, heating system, and CO_2_ control (37 °C, 5% CO_2_). Images were acquired every 10 min using an Andor Neo 5.5 sCMOS camera, a 10× Plan Fluor air objective, and a Lumencor SOLA light engine (phase-contrast for cells, wide-field epifluorescence for beads). At the end of each timelapse, a reference frame was acquired after cell detachment with 200 µL of 10% SDS or Triton X-100.

Fluorescent bead images were registered against the reference frame to calculate displacements. Pre-processing in FIJI included stabilization and illumination correction. Displacement fields were extracted using PIVlab [54] (32×32-pixel windows, 50% overlap). Traction forces were computed with Fourier Transform Traction Cytometry, and intercellular stress fields were reconstructed via Bayesian Inversion Stress Microscopy [55]. Isotropic stress was defined as (*σ*_*xx*_ + *σ*_*yy*_)/2, with positive values denoting tension and negative values compression.

## ACKNOWLEDGEMENTS

We thank Martin Cramer Pedersen, Tianxiang Ma, and Hengdong Lu for the help with PCA and stress analyses. V. S. acknowledges funding from Knut and Alice Wallenberg Foundation (V.S.) via the Wallenberg Centre for Molecular Medicine, Lund; Cancerfonden (19 0445 Pj and 22 2398 Pj Projekt grant). A. D. acknowledges funding from the Novo Nordisk Foundation (grant No. NNF18SA0035142 and NERD grant No. NNF21OC0068687), Villum Fonden (Grant no. 29476), and the European Union (ERC, PhysCoMeT, 101041418). Views and opinions expressed are however those of the authors only and do not necessarily reflect those of the European Union or the European Research Council. Neither the European Union nor the granting authority can be held responsible for them.

**Bibliography**

